# Unbiased detection of CRISPR off-targets *in vivo* using DISCOVER-Seq

**DOI:** 10.1101/469635

**Authors:** Beeke Wienert, Stacia K Wyman, Christopher D Richardson, Charles D Yeh, Pinar Akcakaya, Michelle J Porritt, Michaela Morlock, Jonathan T Vu, Katelynn R Kazane, Hannah L Watry, Luke M Judge, Bruce R Conklin, Marcello Maresca, Jacob E Corn

**Author notes:** the authors contributed equally to this work (Beeke Wienert, Stacia K Wyman).

## Abstract

Genome editing using nucleases such as CRISPR-Cas induces programmable DNA damage at a target genomic site but can also affect off-target sites. Here, we develop a powerful, sensitive assay for the unbiased identification of off-target sites that we term DISCOVER-Seq. This approach takes advantage of the recruitment of endogenous DNA repair factors for genome-wide identification of Cas-induced double-strand breaks. One such factor, MRE11, is recruited precisely to double-strand breaks, enabling molecular characterization of nuclease cut sites with single-base resolution. DISCOVER-Seq detects off-targets in cellular models and *in vivo* upon adenoviral gene editing of mouse livers, paving the way for real-time off-target discovery during therapeutic gene editing. DISCOVER-Seq is furthermore applicable to multiple types of Cas nucleases and provides an unprecedented view of events that precede repair of the affected sites.

## Introduction

CRISPR-Cas (clustered regularly interspaced short palindromic repeats; CRISPR-associated) genome editing holds great promise for therapeutic applications. CRISPR-Cas nucleases make double-strand breaks (DSBs) at the intended target site but can also introduce unwanted mutations at off-target sites within the genome. To bring CRISPR into the clinic, accurate characterization of on- and off-target nuclease activity in any target cell population is critical (*1*).

A variety of *in silico* (*2*–*4*), *in vitro* (*5*–*7*), and cell-based assays (*8*, *9*) have been developed to predict or find CRISPR off-targets. While each of these orthogonal methods have their particular strengths, they also have certain weaknesses. Naïve prediction algorithms are for the most part based on sequence similarity and currently have limited predictive power with very high false-positive rates (*10*). Assays that induce DSBs *in vitro,* such as Digenome-Seq (*5*), CIRCLE-Seq (*6*) and SITE-Seq (*7*), have high sensitivity but dramatically under- or overestimate the number of target sites that are actually modified in cellular models or *in vivo* (*11*). Nuclease concentration within the cell (*7*), delivery method (ribonucleoprotein (RNP) vs. plasmid) (*7*, *12*, *13*) as well as more complex cellular properties such as chromatin accessibility (*14*, *15*) have been shown to significantly affect editing outcomes and are generally missed by *in vitro* off-target assays. Cellular assays, such as GUIDE-Seq (*8*), test nuclease cutting in a cellular context but rely on the integration of an exogenous DNA oligo that is inefficient in primary cells and not applicable *in vivo*. Furthermore, the co-transfection of additional exogenous DNA would not be used in human therapy and may affect overall editing outcomes (*16*). Hybrid methods determine potential editing sites *in vitro* followed by exhaustive testing *in vivo* via amplicon sequencing, but require testing of hundreds to thousands of candidate sites with high false-positive rates (*11*). There remains a pressing need for unbiased methods that characterize sites of genome editing *in situ* in human cells and animal models without requiring extensive pre-treatment or introduction of additional editing reagents beyond those used in gene editing therapies.

Here, we asked whether the recruitment of DNA repair proteins can be developed into a generally applicable assay for the unbiased identification of sites of genome editing. Because the pathways involved in DSB repair are broadly conserved among metazoans, such an approach is potentially useful in a wide variety of contexts. After determining the recruitment kinetics of several repair factors, we developed a powerful assay for the unbiased identification of off-target gene editing in cells and organisms we term <u>D</u>iscovery of *<u>I</u>n <u>S</u>itu* <u>C</u>as <u>O</u>ff-targets and <u>VER</u>ification by <u>Seq</u>uencing (DISCOVER-Seq). This approach involves chromatin immunoprecipitation followed by sequencing (ChIP-Seq) of MRE11, followed by a custom computational pipeline called BLunt END FindER (BLENDER). We find that MRE11 is broadly expressed across tissues and recruited very early to a DSB with such precision that we are able to identify molecular characteristics of nuclease cut sites with single nucleotide resolution. We demonstrate that DISCOVER-Seq reliably detects off-target editing from a variety of guide RNAs, multiple Cas nucleases, in human and mouse cells, and can even determine tissue and locus specificity during adenovirus-mediated *in vivo* gene editing of mice.

## Results

### Distinct binding characteristics of DNA repair proteins at Cas-induced DSBs

While ChIP-Seq for catalytically inactive Cas9 nuclease has been explored to find off-targets (*17*, *18*), this approach is plagued by many false positives because Cas9 binds far more sites than it actually cleaves (*19*, *20*). However, assembly of DNA repair proteins at Cas target sites indicates that DNA damage has occurred, and by identifying DNA sequences that are bound by DNA repair machinery one might theoretically map nuclease-induced sites of DNA damage genome-wide.

Several DNA repair proteins produce distinct microscopic foci at the locations of DSBs induced by radiation or UV light (*21*). However, the molecular behavior of these factors at Cas nuclease-induced break has not been studied in detail. We asked how various repair factors assemble at Cas9-induced DSBs and if we could use this information to identify Cas9 cut sites in an unbiased fashion. We began by performing ChIP-Seq experiments in K562 cells that were nucleofected with a Cas9-RNP targeting *VEGFA* (**Fig. 1A**). We selected a panel of proteins involved in DNA repair of DSBs: Ser193-phosphorylated histone 2AX (γH2AX), XRCC6/Ku70, FANCD2 and the three components of the MRN complex: MRE11, RAD50 and NBS1. We observed that all six DNA repair proteins assembled at the targeted genomic location, but their binding patterns across the site were highly distinctive (**Fig. 1B**). Both γH2AX and FANCD2 produced a broad peak centered at the target site kilobases (kb) to megabases (mb) in length. Chromatin within 1-2 kb of the DSB showed reduced occupancy by γH2AX, consistent with previous reports (*22, 23*). Ku70 and members of the MRN complex showed a defined narrow peak at the cut site, consistent with their known roles in end processing (*24*–*26*) (**Fig. 1C**).

**Figure 1:**
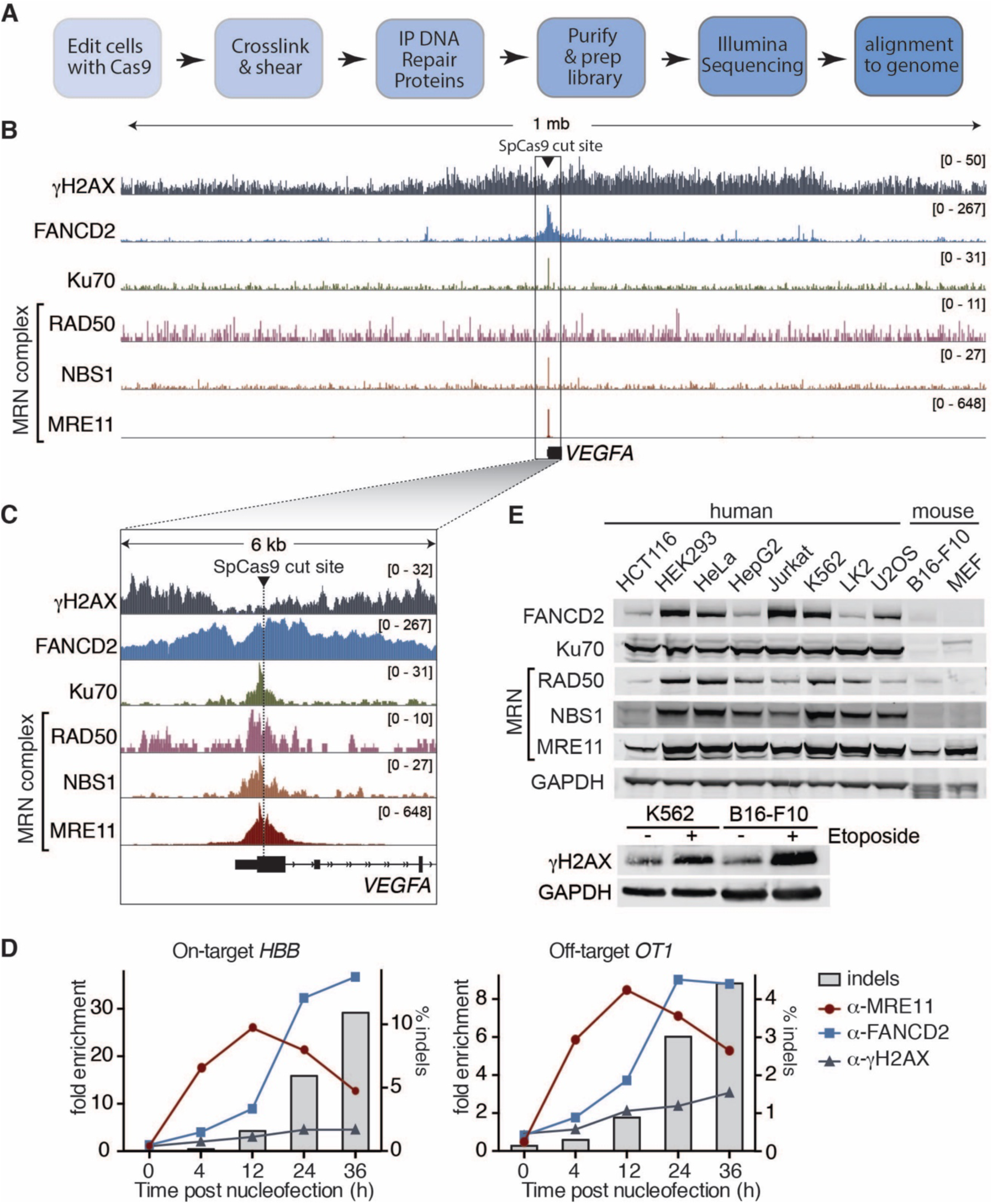
DNA repair proteins assemble at a Cas-induced DSB. **(A)** Schematic of workflow used to determine DNA repair protein binding properties at Cas9-induced DSBs. **(B)** γH2AX, FANCD2, Ku70, NBS1, RAD50, and MRE11 accumulate at SpCas9 cut sites in cells. DNA repair proteins were localized and quantified by ChIP-Seq using the indicated repair proteins in K562 cells and plotting ChIP-Seq reads over a 1Mb window spanning the *VEGFA* cut site. Inset numbers define the height of the y-axis in reads. **(C)** Height and width of ChIP-Seq signals vary widely for DNA repair proteins. Data presented as described in Fig. 1B, except the windowed region is 6kb. The SpCas9 target site is marked with a dotted line. **(D)** DNA repair proteins are present at Cas9-induced breaks with varying kinetics. ChIP-qPCR experiment for MRE11, FANCD2 and γH2AX in K562 cells edited at t=0 with an RNP targeting the *HBB* gene. Cells were harvested for ChIP at indicated time points and DNA extracted. The fold enrichment is the relative binding of each factor at the on-target site (*HBB*) or a known off-target site (*OT1*) normalized to a negative control region (*VEGFA*). The bar graph depicts the amount of indels as determined by amplicon-NGS. Shown is one replicate, a second replicate is shown in Fig. S1. **(E)** Protein levels of DNA repair proteins across a variety of human cell lines from different origins and cross-reactivity of the tested antibody with two murine cell lines. Whole cell lysates from untreated cells were analyzed for FANCD2, Ku70, MRE11, NBS1 and RAD50 abundance by Western Blot. GAPDH abundance is used as loading control. To induce γH2AX phosphorylation, cells were treated with Etoposide for 16h before protein extraction.

### The dynamics of DNA repair proteins at Cas9-induced DSBs

To investigate how the binding of repair factors coincides with the formation of genetic insertions and deletions (indels, the finished product of DNA repair) at the target site, we performed ChIP-quantitative PCR (ChIP-qPCR) time course experiments using K562 cells edited with a Cas9-RNP targeting the *hemoglobin beta* (*HBB*) gene. We monitored both indel formation and binding of γH2AX, FANCD2 and MRE11 to the *HBB* target and a previously described off-target site (*OT1*) (*27*, *28*) (**Fig. 1D, Fig. S1** and **Table S1**). The presence of FANCD2 and γH2AX at the cut site increased over time concurrently with the formation of indels, but MRE11 binding peaked at 12 hours post nucleofection and reduced as indels formed. These data suggest that MRE11 binds to the ends of a DSB but then disassociates, whereas FANCD2 and γH2AX may persist even as indels are forming at a site. MRE11 is therefore a good candidate to detect sites of Cas cleavage, since its action is immediate to the break both in space and time.

For an off-target discovery approach to be broadly useful, it must be feasible in as many cell types and organisms as possible. In this case, it is important that the DNA repair protein of interest is ubiquitously expressed. We therefore tested protein expression levels of DNA repair proteins in a panel of human cell lines from diverse origins. Furthermore, we tested cross-reactivity of the antibodies used for ChIP-Seq between human and mouse, since an ideal off-target discovery method would be equally applicable not just in human cells but also in mice that are commonly used as a model organism for therapeutic editing. Of all proteins tested, only MRE11 showed high expression in all tested cell lines and was also recognized by an antibody that cross-reacts with murine Mre11 (**Fig. 1E**).

### MRE11 ChIP-Seq provides single-nucleotide resolution of Cas-induced DSB

We examined the ChIP-Seq reads associated with each repair protein at the *VEGFA* target site and found that they provide molecular insight into each protein’s role in DNA repair. We classified read pairs that start or end precisely at the cut site (e.g. are “blunt”), indicating binding to the ends of a DSB. We also classified read pairs that “span” the cut site (**Fig. 2A**, left).We found that DNA bound by FANCD2 and γH2AX predominantly span the cut site, though γH2AX occupancy was highest outside the cut site, consistent with prior reports (*22*) (**Fig. 1A**). DNA bound by the MRN complex or Ku70 was primarily blunt at the cut site (**Fig. 2A**, right). These data confirm the roles of Ku70 and the MRN complex as early responders to DSBs after Cas9-induced DNA damage and illustrate that DNA fragments pulled down by MRN/Ku70 ChIP contain unrepaired DSBs (*21*).

**Figure 2:**
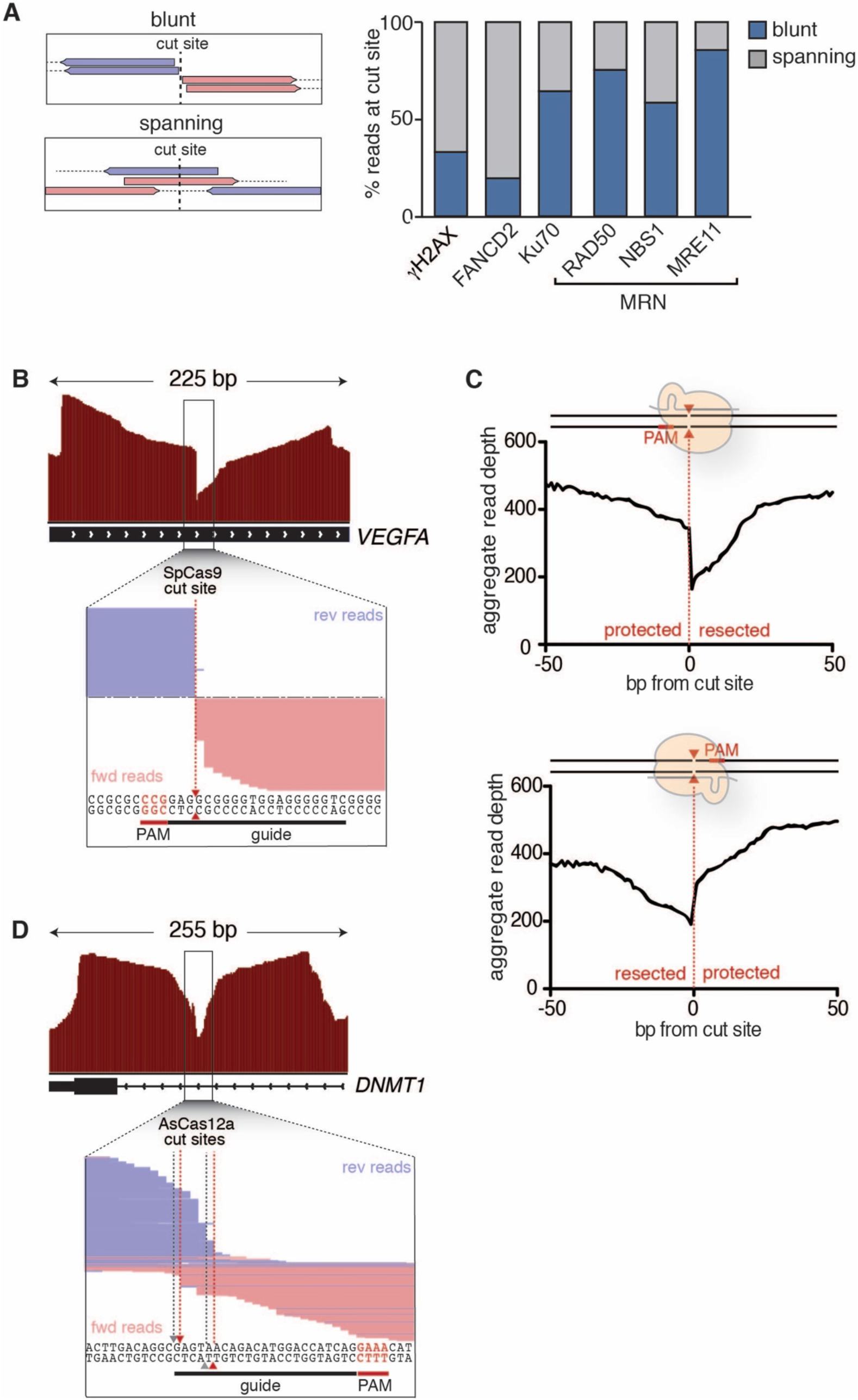
ChIP-Seq molecularly characterizes Cas-induced breaks at single-base resolution. **(A)** Schematic of different types of reads found at the cut site (left) and composition of aligned ChIP-Seq reads at the on-target cut site (*VEGFA*) for each DNA repair factor (right). Shown is the fraction of read pairs that end or start within a 10 bp window around the cut site (blunt) and the fraction of read pairs that are spanning the cut site (spanning). **(B)** MRE11 peaks are asymmetric. MRE11 ChIP-Seq signal is presented as described in **Fig. 1B** over a 225 bp interval spanning the cut site. The windowed region (black box) shows aligned individual Illumina Sequencing reads in forward (pink) or reverse (purple) direction. MRE11 peaks have a distinct shape due to reads ending or starting in the Cas9-induced cut site. Peaks are markedly asymmetric, since all reads perfectly abut one side of the break but many reads are resected from the other side (see **Fig. S2A** for more detail). **(C)** Accumulated read depth of MRE11 ChIP-Seq peaks in K562 cells edited with SpCas9-RNPs targeting *VEGFA.* Reads are accumulated over multiple on- and off-target sites, binned into whether the gRNA binds the sense strand (eight sites) or the gRNA binds the antisense strand (eight sites). Reads are blunt against the PAM-containing end of the DNA, but are both blunt and resected on the non-PAM end. **(D)** MRE11 ChIP-Seq in K562 cells edited with an AsCas12a-RNP targeting *DNMT1*. Enlarged is a region around the AsCas12a cut site showing the composition of aligned reads. The read distribution matches the activity of AsCas12a to produce a 5’ overhang. A predicted 4 nucleotide AsCas12a cut site is marked in grey and a 5 nucleotide alternative cut site is shown in red.

Strikingly, we found that ∼90% of the reads in MRE11 peaks were blunt at the Cas9-induced cut site. Paired-end 100-mer Illumina sequencing reads from MRE11 ChIP-Seq begin randomly on one side of the break, due to fragmentation during sonication, and then end exactly at the Cas9 cut site (**Fig. S2A**). While the insert size between the two paired ends varies, sequencing always starts at the cut end, and thus alignment of paired reads causes a stack of 100-mers to each end of the cut site causing a characteristic shape of MRE11 ChIP-Seq peaks (**Fig. S2A**). The blunt side of MRE11 binding to a Cas9-induced DSB is in fact so precise that the exact nucleotide corresponding to the *in vitro* identified Cas9 nuclease site is identifiable by examining an MRE11 peak (**Fig. 2B**). Precise identification of a Cas9 cut site was possible for both the on-target site of a *VEGFA* targeting guide RNA (gRNA), as well as 32 known off-target sites (*8*) (**Fig. S2B**). To the best of our knowledge, this is the first molecular observation of Cas9’s *in vitro* nuclease activity in a cellular context.

While the MRE11 peak at a Cas9 DSB is blunt on the PAM side of the *VEGFA* on-target site, it is much more resected on the non-PAM side and hence asymmetric (**Fig. 2B, Fig. S2A-B**). Since we previously observed that the non-PAM side of a Cas9 DSB is released first *in vitro* (*29*), we asked if MRE11 peak asymmetry is predictable based on the location of the PAM. We aggregated MRE11 peaks from the *VEGFA* on-target and 15 known off-targets and binned them based on whether the PAM is oriented in the sense or anti-sense direction (**Fig. 2C**). Strikingly, the asymmetry of a Cas9-induced MRE11 peak depends upon on the PAM orientation, such that the blunt reads accumulate on the PAM side and the resected reads accumulate on the non-PAM side (**Fig. 2C**). These data suggest that either the PAM side of a Cas9-induced DSB is protected against early resection by cellular nucleases (*30*) or that cutting by Cas9 itself is not as precise on the non-PAM side (*31*), both of which are equally consistent with *in vitro* data. Our observation of asymmetric resection data in living cells is also consistent with *in vitro* and cellular data that Cas9 holds on more tightly to the PAM side and protects it from enzymatic modification (**Fig. S2C**) (*29*). It may be that the asymmetric nature of Cas9-induced break influences the highly stereotyped indel spectra reported for Cas9 editing (*32*–*34*).

While SpCas9 makes relatively blunt-ended cuts (*31*), other nucleases such as Cas12a (also known as Cpf1) produce staggered DSBs with a four nucleotide 5’-overhang (*35*). However, these digestion patterns are based on *in vitro* data and it has so far not been possible to directly visualize Cas12a nuclease activity in cells. We asked if MRE11 binding could be used to reveal Cas12a activity during editing by nucleofecting an *Acidaminococcus sp.* Cas12a (AsCas12a) RNP targeting the *DNMT1* gene into K562 cells and performed ChIP-Seq for MRE11 (**Fig. 2D**). Unlike SpCas9 MRE11 peaks, the AsCas12a MRE11 peak was quite symmetric around the predicted cut sites. Furthermore, the reads at the targeted site overlapped one another but were blunt in characteristic locations, consistent with the production of an overhang. However, the cellular pattern of AsCas12a was not strictly a defined 4 nucleotide overhang as suggested *in vitro*, but instead is ambiguous between 4-6 nucleotides. There is some evidence of mixed cleavage by AsCas12a from *in vitro* studies (*35*) and our results indicate that the molecular site of nuclease activity by AsCas12a in cells is not as simple as previously assumed. Taken together, our results show that MRE11 ChIP-Seq can characterize nuclease-induced DSBs at a molecular level for multiple types of Cas nucleases.

### DISCOVER-Seq detects off-targets in human cells

While performing editing with the *VEGFA* targeting RNP, we manually identified MRE11 peaks at known off-target sites for this gRNA. We next asked if we could develop MRE11 ChIP-Seq into an unbiased method to discover CRISPR-Cas9 off-targets. We term this approach <u>D</u>iscovery of *<u>I</u>n <u>S</u>itu* <u>C</u>as <u>O</u>ff-targets and <u>Ver</u>ification by ChIP-Seq (DISCOVER-Seq). The general workflow of DISCOVER-Seq is depicted in **Fig. 3A** and involves MRE11 ChIP-Seq followed by a bioinformatic pipeline called BLENDER (BLunt ENd finDER) to identify the characteristic read signatures of Cas-induced breaks. BLENDER traverses genome-wide ChIP-Seq data, locating stacks of reads and scoring each site based on the summed read ends within a window around the cut site. BLENDER takes into account the unique features of MRE11 peaks, has several options to optionally black-list known ChIP-Seq artifacts, references untreated control samples for additional sensitivity, and filters based on protospacer identity for reduced false positives to thus call even rare off-target events with great accuracy (**Fig. S3**).

**Figure 3:**
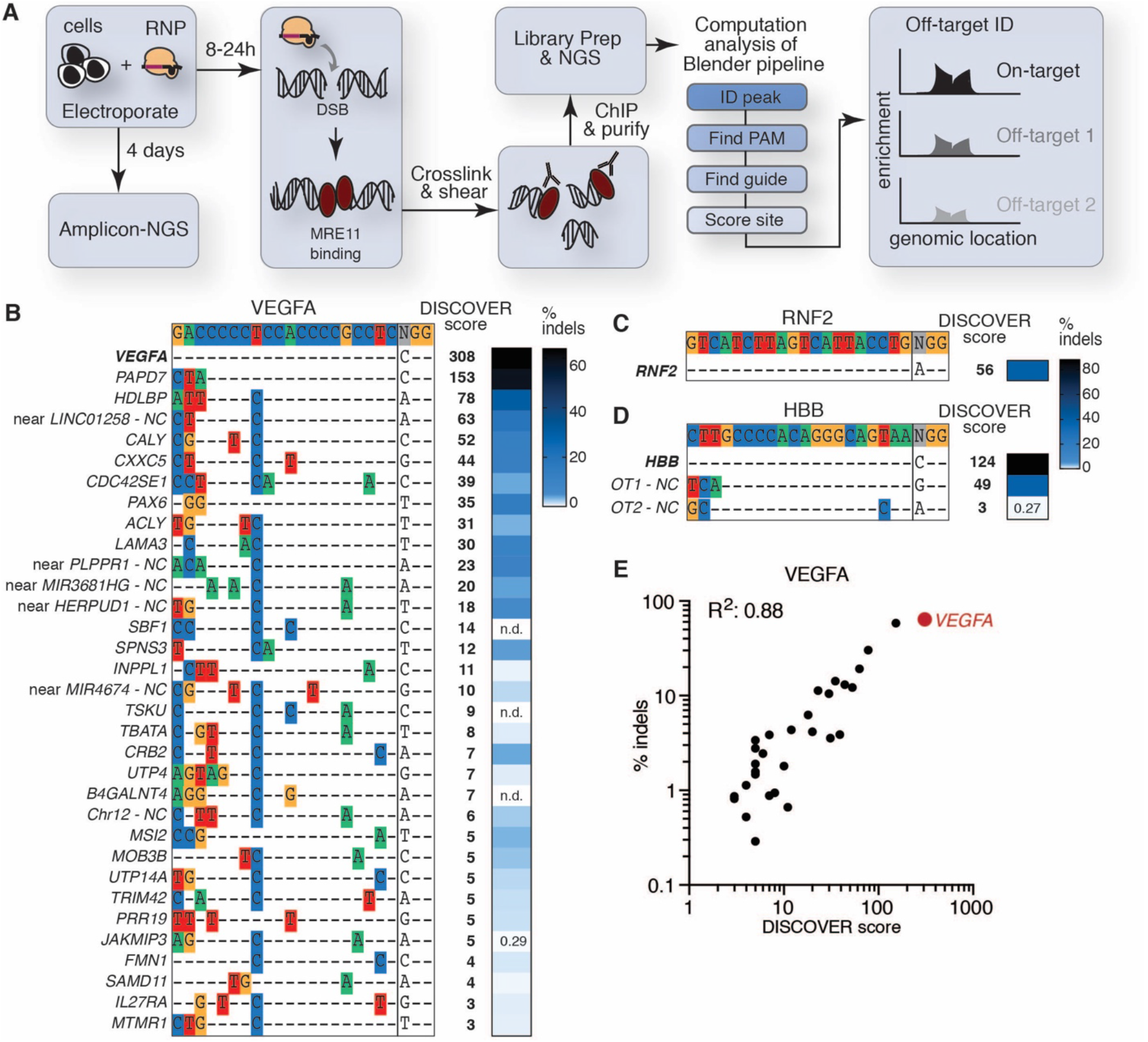
Unbiased off-target discovery in human cells using DISCOVER-Seq. **(A)** Schematic showing the general workflow of DISCOVER-Seq. The first part of DISCOVER-Seq consists of genome editing of cells followed by MRE11 ChIP-Seq at a relatively short time point to capture Cas-induced DSBs. The data is then input to a custom bioinformatics pipeline (BLENDER) to determine the genomic location of off-target sites (see Fig. S3 for more detail). **(B)** Sequences of off-targets identified with DISCOVER-Seq for *VEGFA* site 2 in K562 cells. The on-target sequence and site is shown on the top and discovered off-target cleavage sites are beneath. Any mismatches to the on-target sequence are highlighted in color. DISCOVER scores and indel frequencies compared to unedited cells as determined by amplicon-NGS are shown to the right for each site. DISCOVER-Seq finds *bona fide* off-target sites verifiable with amplicon-NGS and does not report false positive sites. NC: non-coding. n.d.: not determined due to PCR difficulties. **(C)** and **(D)** Sequences, DISCOVER scores and indel frequencies of off-targets identified with DISCOVER-Seq for *RNF2* and *HBB* targeting gRNAs in K562 cells. **(E)** DISCOVER scores and indel frequencies are highly correlated. Relationship between *VEGFA* off-target indel frequencies and DISCOVER score for each off-target. *VEGFA* on-target site is highlighted in red.

We tested DISCOVER-Seq using the well-characterized, promiscuous ‘VEGFA site 2’ gRNA in K562 cells (*8*). Unbiased DISCOVER-Seq identified 32 off-target sites for *VEGFA_site2*, which we validated individually by amplicon next-generation sequencing (amplicon-NGS) four days after nucleofection (**Fig. 3B** and **Table S1**). All off-targets that were identified by DISCOVER-Seq were sites that have previously been described by GUIDE-Seq and were amplicon-NGS validated with indel rates above background, with the least frequent site containing 0.29% indels (**Fig. 3B**).

We also asked the converse, if there are major off-targets missed by DISCOVER-Seq but found by GUIDE-Seq. We performed amplicon-NGS on all putative off-targets previously identified with more than more than 100 GUIDE-Seq reads, but found that none of these sites had indel rates above 1.2% (**Table S1**).

We further tested DISCOVER-Seq on a very specific guide targeting *RNF2* with no known off-targets, and a semi-specific guide targeting *HBB* with two known off-targets. Again, we validated all off-targets found with DISCOVER-Seq by amplicon-NGS (**Fig. 3C-D**). All sites identified with DISCOVER-Seq showed indels above background and were thus *bona fide* off-targets. For the *RNF2* guide, DISCOVER-Seq found no sites other than the on-target site, confirming its specificity (*8*) (**Fig. 3C**). For the *HBB* guide, DISCOVER-Seq successfully identified the two previously described off-target sites (*27*, *28*) (**Fig. 3D**). Overall, these data show that DISCOVER-Seq is very capable of unbiased *in situ* off-target detection in human cells. DISCOVER-Seq can distinguish a specific gRNA from an unspecific gRNA and sensitively finds *bona fide* off-targets with no false positives (**Fig. 3E**). Furthermore, indel frequencies and DISCOVER scores were highly correlated suggesting that DISCOVER scores can be used to predict off-target mutation rates (**Fig. 3E**).

### Off-target discovery in murine cells

As the MRE11 antibody used for DISCOVER-Seq is cross-reactive with the mouse protein, it can theoretically be used in a murine setting without modification of the protocol. Thus, we tested off-target discovery with previously characterized promiscuous and highly specific gRNAs in murine cells. We edited the mouse skin melanoma cell line B16-F10 with two different Cas9-RNPs targeting the *Pcsk9* gene and performed DISCOVER-Seq on these cells. The promiscuous *Pcsk9* gRNA, “gP”, has many closely matched sites in the mouse genome and thus many off-targets, while the specific gRNA, “gM”, and has no off-targets (*11*). For gP edited cells, DISCOVER-Seq found 44 off-target sites in addition to the on-target *Pcsk9* site (**Fig. 4A** and **Fig. S4A**). In this experiment, we found that DISCOVER-Seq was equally capable of identifying off-targets with the canonical SpCas9 PAM (NGG) and a known non-canonical PAM (NAG). For gM edited cells, DISCOVER-Seq exclusively found the on-target site (**Fig. 4B**). To validate the DISCOVER-Seq identified sites, we performed individual amplicon-NGS to determine indel frequencies after 4 days. We also performed amplicon-NGS at additional sites previously reported for the gP guide (*11*) (**Fig. 4A-B** and **Table S2**). All seven previously reported sites not identified by DISCOVER-Seq were very inefficiently edited, with indel frequencies below 0.43 % and four of them had indels below 0.1 % (**Table S2**). DISCOVER scores and indel rates were correlated for both the gP and gP+G guide RNAs (**Fig. S4A**). We found that sites identified and validated with DISCOVER-Seq had indel rates ranging from 0.1% to 66.1%, showing that DISCOVER-Seq is applicable for off-target detection in murine cell systems.

**Figure 4:**
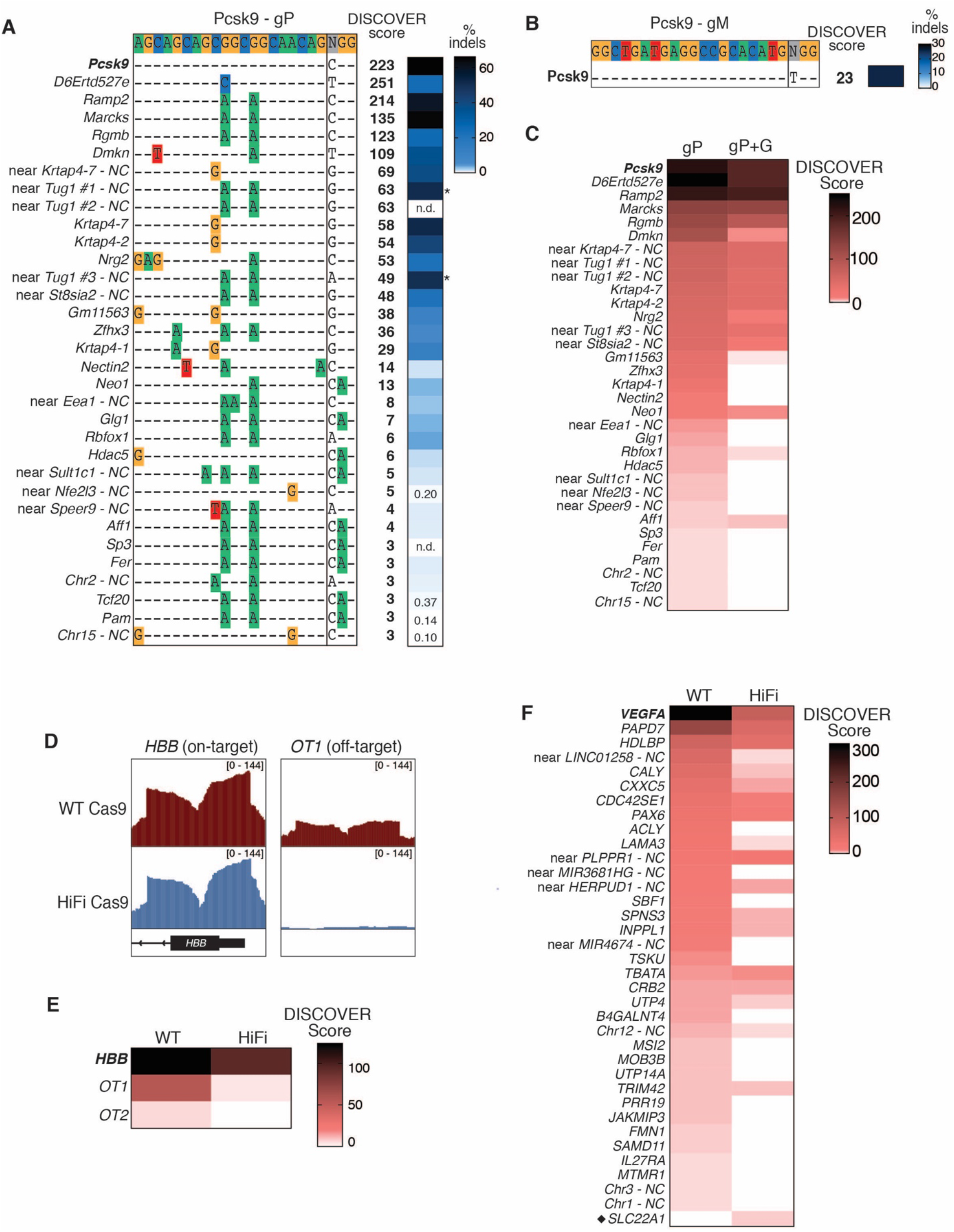
DISCOVER-Seq is broadly applicable. **(A)** Sequences of off-targets for the *Pcsk9* gP gRNA identified with DISCOVER-Seq in murine B16-F10 cells. The on-target sequence and site is shown on the top and discovered off-target cleavage sites are beneath. Any mismatches to the on-target sequence are highlighted in color. DISCOVER scores and indel frequencies compared to unedited cells as determined by amplicon-NGS are shown to the right for each site. Indel frequencies marked with an asterisk were not reliable by amplicon-NGS and were instead determined by Sanger sequencing and ICE analysis. NC: non-coding; n.d.: not determined. **(B)** Sequence, DISCOVER score, and indel frequency of the single target for the *Pcsk9* gM gRNA identified with DISCOVER-Seq in murine B16-F10 cells. **(C)** DISCOVER scores for off-targets of the *Pcsk9* gP gRNA and a *Pcsk9* gP gRNA with an additional 5’ G added to the protospacer (gP+G) in B16-F10 cells. The addition of the 5’ G, commonly added for efficient *in vitro* transcription and plasmid or viral transcription, has a large effect on the number of off-targets. **(D)** MRE11 ChIP-Seq peaks for K562 cells edited with WT Cas9 or HiFi Cas9 and *HBB* gRNA at the on-target (*HBB*) and a known off-target site (*OT1*). **(E)** DISCOVER scores for off-targets from K562 cells edited with WT Cas9- or HiFi Cas9-RNPs targeting *HBB*. **(F)** DISCOVER scores for off-targets from K562 cells edited with WT Cas9- or HiFi Cas9-RNPs targeting *VEGFA* (*VEGFA* site 2). A black diamond indicates an off-target that was specific for the sample edited with HiFi Cas9 and has not been previously described.

### Common additions to gRNAs alter off-target profiles

Many research laboratories generate their gRNAs by *in vitro* transcription (in the case of RNP editing) or transduce cells with plasmids or virus encoding gRNA. To ensure efficient transcription from the T7 or U6 promoter, the protospacer must either begin with a 5’ guanine (G), or an extra 5’ G must be added resulting in a 21-nt protospacer. Synthetic gRNAs do not require an extra 5’ G and can begin with any nucleotide. While the 5’ G addition is relatively innocuous in terms of on-target editing efficiencies, there is little data on how it affects off-target editing. Yet without such knowledge, comparisons of off-target profiles between guides that contain an extra 5’ G and those that do not are potentially flawed.

The promiscuous *Pcsk9* gP guide we used previously does not have a G in the first position but has previously been used in mice via viral transduction (*11*) and thus was characterized with an additional 5’ G. We used DISCOVER-Seq to ask if adding an extra 5’ G to *Pcsk9* gP changes the off-target profile of the gRNA. We edited B16-F10 cells with Cas9 and a gP guide containing an extra 5’ G (gP+G) and performed DISCOVER-Seq. Comparing DISCOVER scores of gP and gP+G, we found that the additional G markedly reduced off-target frequencies (**Fig. 4C** and **Fig. S4B**). However, this reduction was not consistent across sites, and some off-target loci were still detected with high frequency. Amplicon-NGS confirmed that on-target indel frequencies were comparable between gP and gP+G edited cells, but off-target mutation rates differed substantially (**Fig. S4C** and **Table S2**). Thus, DISCOVER-Seq can not only distinguish high-fidelity gRNAs from promiscuous gRNAs, but can also be used to characterize off-target profiles generated by gRNAs with alternative protospacer lengths and modifications.

### Cas9 cutting dynamics at on- and off-target sites

The kinetics of Cas9 cleavage and binding are well-studied *in vitro* and the amount of mismatches of a protospacer to an on- or off-target site can dramatically affect an RNP’s on- and off-rate (*36*). However, relatively little is known about dynamics of on- and off-target cleavage by Cas9 in a cellular context. Because DISCOVER-Seq measures DSBs rather than finished repair outcomes (indels), we used it to ask if off-target sites are cut simultaneously with the on-target site. We performed a DISCOVER-Seq time-course experiment in B16-F10 cells edited with Cas9 and the promiscuous gP+G guide, analyzing samples 8h, 12h, 24h and 36h post nucleofection (**Fig. S4D**). We observed that DSBs at the on-target site were most prominent at the 12h time point, supporting our observations from the ChIP-qPCR time course in K562 cells (
**Fig. 1D** and **Fig. S1**). We found that off-target cleavage dynamics were generally comparable to those at the on-target site and that earlier time points allow a more sensitive detection of off-targets (**Fig. S4D**).

### DISCOVER-Seq characterizes off-target profiles of Cas9 variants

Several high-fidelity Cas9 variants have been developed that reduce off-target editing (*37*–*39*). Since DISCOVER-Seq can distinguish high-specificity gRNAs from promiscuous gRNAs, we asked if this approach can also distinguish a high-fidelity Cas9 mutant from the less fidelitous wild type (WT) Cas9. We edited K562 cells with RNPs targeting the previously mentioned *HBB* locus that has two known off-target sites. We formed these RNPs using WT Cas9 and the HiFi Cas9 R691A mutant (*38*) and compared the DISCOVER-Seq profiles of each experiment. Cells edited with WT Cas9 showed prominent MRE11 peaks at the on-target site in *HBB* and off-target site *OT1*, with a smaller peak at the less frequently edited *OT2* site. Cells edited with HiFi Cas9 showed comparable MRE11 binding at *HBB,* but a dramatically decreased binding of MRE11 to *OT1* and undetectable binding to *OT2* (**Fig. 4D**). Amplicon-NGS verified that WT Cas9 produces indels at *HBB, OT1,* and *OT2*, while HiFi-Cas9 primarily edits *HBB* with greatly reduced indels at *OT1* and none at *OT2* (**Fig. 4E** and **Fig. S5A** and **Table S1**). To investigate a larger off-target panel we also tested HiFi-Cas9 and DISCOVER-Seq with the promiscuous VEGFA site 2 gRNA in K562 cells. Again, DISCOVER-Seq detected fewer off-targets with the HiFi Cas9 and this was validated with amplicon-NGS (**Fig. 4F** and **Fig. S5B-C** and **Table S1**). Surprisingly, DISCOVER-Seq identified an additional off-target with the HiFi Cas9 that was not found previously by GUIDE-Seq and is not observed with wild-type Cas9 (*8*) (**Fig. 4F**). This new off-target was validated by amplicon-NGS and showed 0.75% indels above background. These data illustrate that high-fidelity Cas9 variants can have unanticipated off-target profiles that can be found using DISCOVER-Seq.

### Off-target detection in patient-derived stem cells using DISCOVER-Seq

We have so far shown that DISCOVER-Seq works robustly in immortalized mammalian cells. Stem cells, such as induced pluripotent stem cells (iPSCs), are much more sensitive and difficult to culture and transfect, and so are not amenable to methods such as GUIDE-Seq. As DISCOVER-Seq requires nothing but the editing reagents to be introduced into cells, off-target detection should be possible even in patient-derived iPSCs. We derived iPSCs from a patient suffering from Charcot-Marie-Tooth disease (CMT) – one of the most common inherited neurological disorders. The patient carries a heterozygous missense mutation (P182L) in *heat-shock protein B1 (HSPB1)* resulting in a dominant negative neuropathic phenotype with a loss of motor neuron function (*40*). Mice heterozygous for HSPB1 show no disease phenotype, so an attractive therapeutic strategy is to knock out the disease allele without affecting the intact copy of *HSPB1*, (*41*, *42*). For this strategy to move into the clinic, it is important to characterize potential missense-targeting editing reagents in detail, ideally in a patient-specific manner.

We edited iPSCs from the CMT patient with HiFi Cas9-RNPs targeting either the *HSPB1* wild-type (WT) or mutant (Mut) allele and subsequently performed DISCOVER-Seq on both sets of edited cells to identify patient-specific and allele-specific off-targets for each gRNA (**Fig. 5A-B**). For the WT gRNA, we detected the on-target site *HSPB1* and one major off-target in a non-coding region that contains an exact duplication of HSPB1 exon 3. Both targets were validated by amplicon-NGS after 4 days (**Fig. 5C**, top, and **Table S3**). DISCOVER-Seq on patient iPSCs edited with the Mut gRNA exclusively identified the on-target site *HSPB1,* but not the noncoding off-target site that contains a protospacer mutation relative to the Mut gRNA (**Fig 5C**, bottom, and **Table S3**). These data suggest that the Mut gRNA is highly specific for the disease allele.

**Figure 5:**
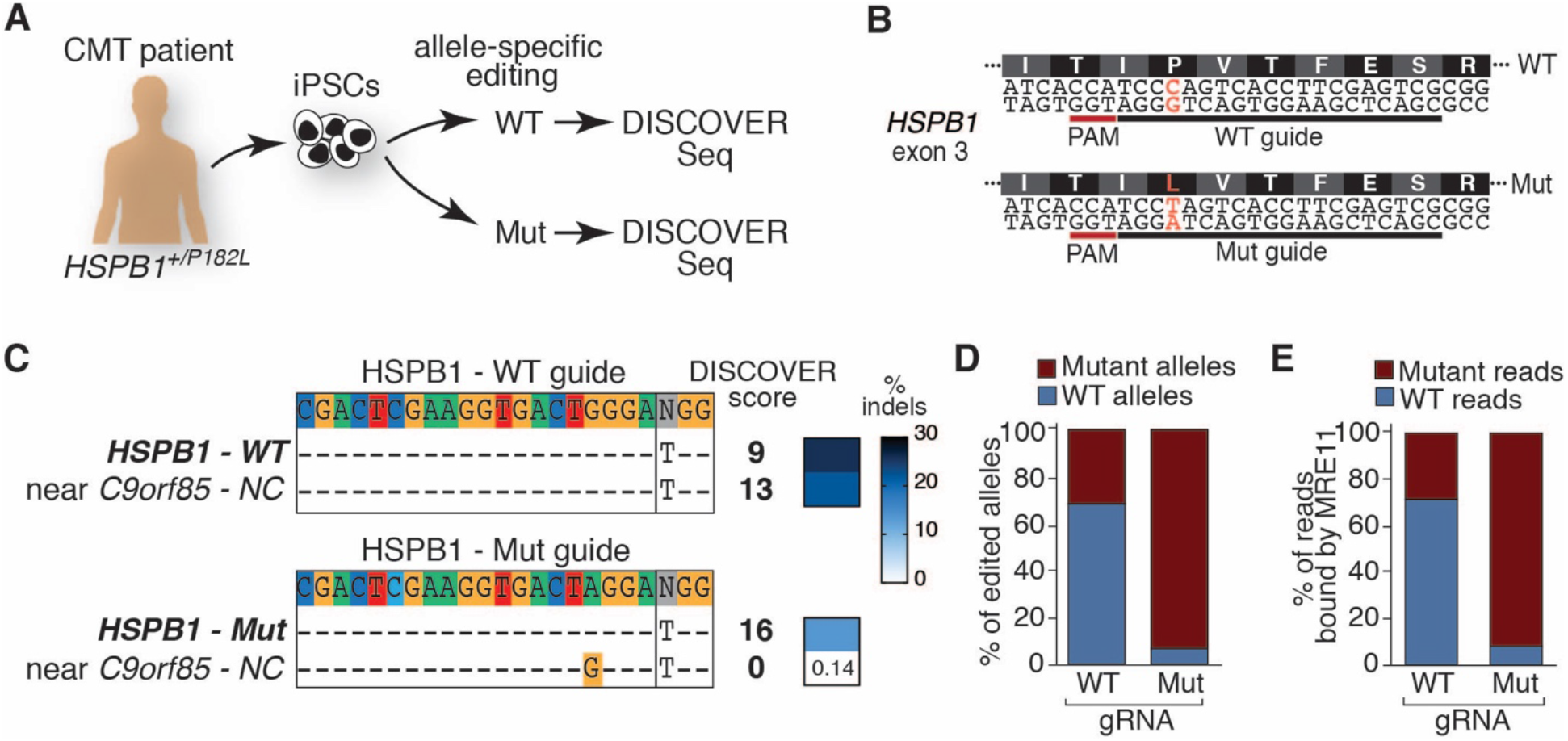
DISCOVER-Seq off-target discovery in patient-derived pluripotent stem cells. **(A)** Schematic showing experimental workflow. iPSCs were derived from a Charcot-Marie-Tooth (CMT) patient heterozygous for the disease mutation P182L in the *HSPB1* gene. iPSCs were edited with HiFi Cas9 and two different gRNAs targeting either the WT or the mutant allele as shown in **(B).** Off-targets from each gRNA were analyzed using DISCOVER-Seq. **(B)** Schematic of alleles in patient iPSCs and targeting gRNAs. **(C)** Sequences, DISCOVER scores, and indel frequencies of off-targets for *HSPB1* WT and *HSPB1* mutant (Mut) gRNAs in heterozygous patient iPSCs. The WT gRNA edits both *HSPB1* as well as a perfectly matching site in a noncoding region, while the Mut gRNA only edits *HSPB1*. **(D)** Allele-specificity of WT and Mut gRNAs was determined by amplicon-NGS. Shown is the fraction of WT and mutant alleles that contain indels after editing with each respective gRNA. Editing with WT gRNA resulted in substantial editing of both WT and mutant alleles while the Mut gRNA almost exclusively targeted the mutant allele. **(E)** Allele-specific editing efficiency correlates with MRE11 recruitment to *HSPB1* cut sites. Shown is the fraction of MRE11 ChIP-Seq reads at the *HSPB1* site that contains either WT or the Mut sequence.

Because indels introduced to the *HSPB1* locus cover the heterozygous patient mutation, simple amplicon-NGS cannot distinguish editing of the WT or Mut allele. Hence, the indel rates reported in **Fig. 5C** are a summary of editing events on both alleles. We therefore used allele drop-out rates to investigate whether editing was allele specific by examining reads that still contained the intact WT or Mut allele in each of the edited samples. In a control sample of heterozygous patient iPSCs that was edited with a non-targeting gRNA, intact WT and Mut reads accounted for approximately 50% of total aligned reads. By subtracting the intact WT or Mut alleles in the edited samples from the observed 50% we determined editing frequencies on each allele (**Fig. 5D**).

We found that cells edited with the WT gRNA showed a substantial reduction of both WT and Mut alleles, suggesting that both alleles were edited to some extent and that the WT gRNA cannot distinguish the two alleles. In contrast, cells edited with the Mut gRNA showed mainly drop-out of Mut alleles but left most WT alleles intact. We then re-purposed DISCOVER-Seq to give us more detailed information about allele-specificity of the WT and Mut gRNAs by examining the sequence of the MRE11 ChIP-Seq reads at the target site. We found that in heterozygous iPSCs edited with the WT gRNA, the DNA bound by MRE11 contained WT sequence but also ∼30% of the G>A mutation. By contrast, in heterozygous iPSCs edited with the Mut gRNA, 92% of DNA bound by MRE11 contained the G>A mutation. (**Fig. 5E**). Overall, these data indicate that the patient-specific *HSPB1* gRNA described here is specific for the mutant allele while sparing the WT allele, and that DISCOVER-Seq can be used to investigate both genomic and allelic off-target sites in patient-specific genomes in order to characterize editing reagents for personalized therapy.

### DISCOVER-Seq measures CRISPR specificity *in vivo* during viral-mediated editing of mice

Characterizing CRISPR-Cas off-targets after *in vivo* gene editing presents a major challenge but is an important milestone for clinical translation of therapeutic editing. Current approaches to test *in vivo* gRNA specificity rely on *in vitro* treatment of isolated genomic DNA followed by exhaustive amplicon-NGS testing of hundreds to thousands of potential off-targets (*11*). While this approach can distinguish promiscuous gRNAs from fidelitous gRNAs, it involves a great deal of experimental effort with many false positives.

We asked if DISCOVER-Seq can be applied to detect off-targets in mouse tissues after *in vivo* viral-mediated gene editing. We first used Western blotting to find that Mre11 expression is variable but broad across multiple mouse tissues and is indeed more widely expressed than other repair proteins such as Fancd2 (**Fig. S6A**).

*In vivo* editing outcomes for the gP+G promiscuous guide targeting *Pcsk9* have been characterized in detail by ‘verification of *in vivo* off-targets’ (VIVO), and so this gRNA represents a good test bed for *in vivo* DISCOVER-Seq (*11*). We used adenoviral infection to deliver gP and WT Cas9 to mice, plus a negative control cohort encoding GFP and Cas9 (**Fig. 6A**). To note, a 5′G is added upon U6 promoter-driven transcription of gP that results in gP expression *in vivo* as gP+G, therefore we denote it here as gP+G. Two mice were sacrificed at each 24h, 36h and 48h timepoint post viral infection and DISCOVER-Seq was performed on the target liver tissue as well as lung tissue that is not targeted by the adenovirus (*11*). As 48h post infection is too early to reliably detect indels *in vivo* (**Fig. S6B**), we used edited livers from three mice that were sacrificed 4 days after infection to determine indel frequencies (**Fig. 6A**).

**Fig. 6:**
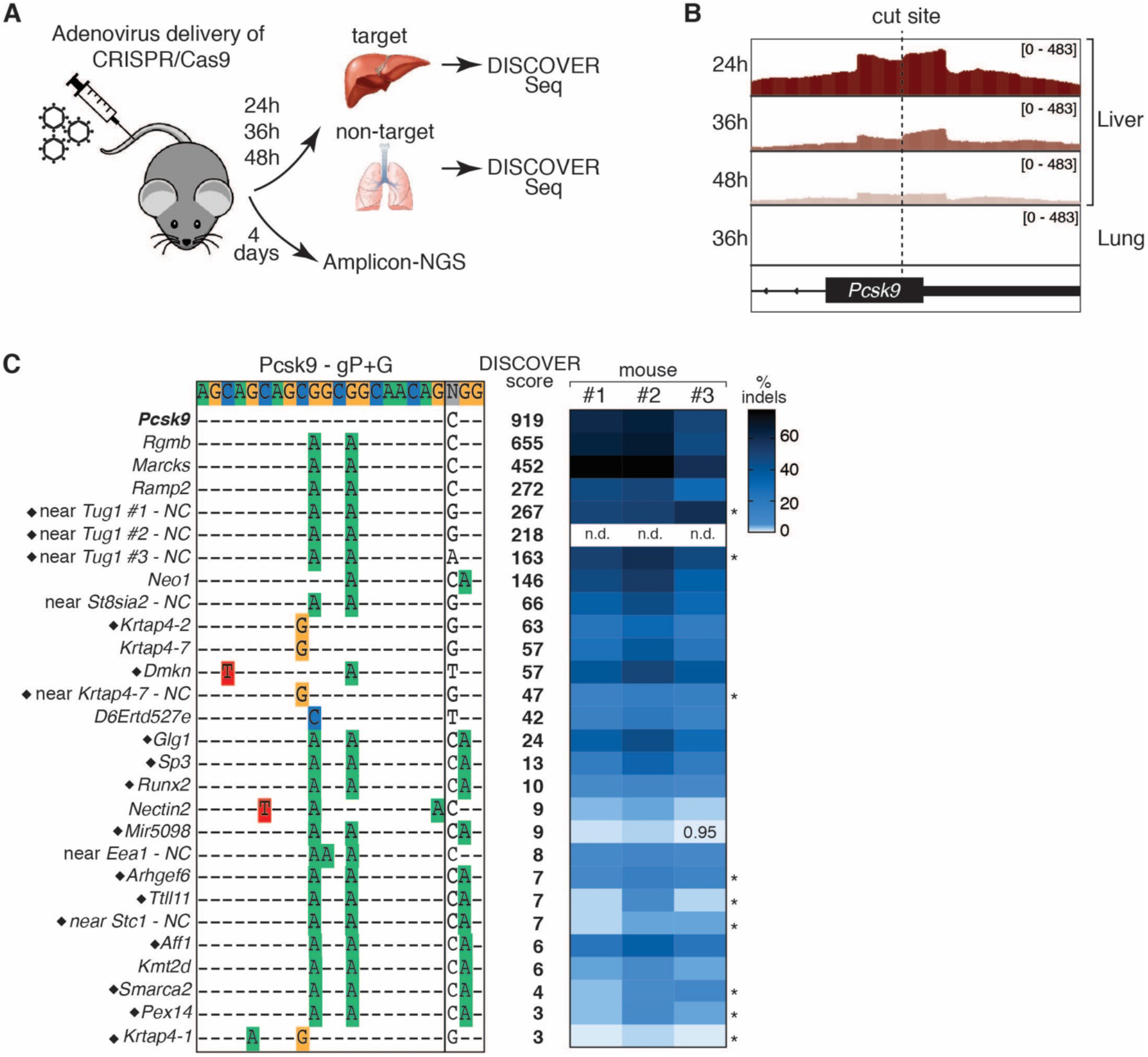
DISCOVER-Seq identifies genome editing specificity at the genetic and tissue level *in vivo*. **(A)** Schematic illustrating the DISCOVER-Seq workflow for *in vivo* gene editing. Multiple mice were injected with hepatotrophic adenovirus encoding either Cas9 and a targeting gRNA (*Pcsk9* gP+G) or a negative control virus encoding Cas9 and GFP. On-target tissues (liver) and off-target tissues (lung) were harvested 24, 36 and 48h after injection and DISCOVER-Seq was performed on the tissues. Livers from injected mice after 4 days were used for amplicon-NGS. **(B)** DISCOVER-Seq accurately distinguishes editing in target and non-target tissues. Mre11 ChIP-Seq signals at the *Pcsk9* on-target site in mouse livers at different time points (top) and lung (bottom). No Mre11 binding was detected at the on-target site in the off-target tissue. Shown are results from three mice at various timepoints, with three additional mice in **Fig. S6B**. **(C)** DISCOVER-Seq identifies sites of on- and off-target editing *in vivo*. Unbiased off-targets identified by DISCOVER-Seq for *Pcsk9* gP+G in mouse livers. The on-target sequence and site is shown on the top and identified off-target sites are beneath. Mismatches to the on-target sequence are highlighted in color. DISCOVER scores and indel frequencies compared to control mice after 4 days as determined by amplicon-NGS or Sanger sequencing and ICE analysis (noted with *) are shown to the right for each site. DISCOVER results from pooled sequencing file of all mice are shown. Black diamonds mark sites that were not previously characterized by VIVO (*11*). NC: non-coding; n.d.: not determined.

Using DISCOVER-Seq, we found that cleavage at the on-target *Pcsk9* site in liver was strongest 24 hours post infection and then declined over time (**Fig. 6B** and **Fig. S6C**). Amplicon-NGS showed between 49.3 and 64.4% indels for the on-target *Pcsk9.* DISCOVER-Seq confirmed tissue-specificity of the adenoviral-mediated editing, measuring no Pcsk9 cleavage in lung tissue, even though Mre11 is well-expressed in the lung (**Fig. 6B** and **Fig. S6A**). We confirmed by amplicon-NGS that no indels were present in the lung tissue (0.042%). Hence, DISCOVER-Seq is capable of establishing tissue specificity during *in vivo* viral genome editing.

To account for potential differences in editing and ChIP-Seq timing between animals, we bioinformatically pooled the DISCOVER-Seq reads from the edited mice and analyzed the pooled reads using BLENDER. To normalize for higher coverage with multiple samples, we adjusted our filtering scheme accordingly to limit false positives (**Fig. S3**). DISCOVER-Seq identified 36 off-target sites in the liver samples and we followed up on 27 of them (**Fig. 6C** and **Table S4**). Amplicon-NGS and Sanger sequencing confirmed that all 27 off-targets that we tested were edited, with indel frequencies ranging from 0.9% to 78.1%. This illustrates that DISCOVER-Seq exclusively identifies *bona-fide* off-targets (**Fig. 6C** and **Table S4**). Low frequency off-targets (<0.8% indels) that were determined as significant by VIVO could not be detected by DISCOVER-Seq. However, 17 of the *bona-fide* off-target sites found by DISCOVER-Seq and validated by amplicon NGS (indel frequencies ranging from 0.95 to 51%) were not previously characterized by VIVO. While CIRCLE-Seq did identify these sites, their CIRCLE-Seq score was relatively low and they could have been missed among the noise of the >3000 sites identified by CIRCLE-Seq. By contrast, DISCOVER-Seq unbiased detection found only true off-targets during viral-mediated *in vivo* gene editing in a single experimental workflow.

## Discussion

DISCOVER-Seq is a universal approach for the unbiased detection of genome editing off-targets that is applicable in cell lines, primary cells, and tissues edited with multiple delivery systems in human and mouse. We find that ChIP-Seq of DNA repair proteins can spatially and temporally characterize Cas-induced DNA damage on a molecular level. In particular, MRE11 ChIP-Seq provides valuable insights into SpCas9 and AsCas12a cleavage patterns and dynamics. We used these data to characterize Cas nuclease activity in living cells. Our data reveal that the PAM side of a Cas9-induced DSB is more protected from resection. Since Cas9 stays tightly bound to DNA *in vitro* (*29*, *30*, *43*, *44*) and *in vivo* (*19*, *20*) even after a DSB has occurred, we favor a model in which the Cas9 protein itself protects the DNA ends on the PAM side from being accessed by other nucleases which could influence repair of the break. Cas9 can itself make non-blunt cuts on the PAM-distal side, but the degree of resection we observe in the MRE11 ChIP-Seq is far greater than the few bases of imprecision observed for *in vitro* Cas9 cutting (*31*). Interestingly, we found no asymmetry after AsCas12a editing, although more data from multiple different Cas12a gRNAs will be needed to support this hypothesis.

DISCOVER-Seq gives insight into the dynamics of Cas9 cutting at on- and off-target sites after RNP editing in cells and after adenoviral editing in mice. This is in contrast to previous cellular kinetics experiments (*45*), which mainly examined the products of repair. We found that DSBs and MRE11 binding occur very early on after RNP editing (4-12h post nucleofection) and as early as 24h after viral infection in tissues. *In vitro* Cas9 resides significantly longer at the on-target site than at off-target sites, resulting in more efficient cleavage (*19*, *20*, *36*). However, our ability to study cleavage products in cells suggest that in this context the on-target and off-target cutting dynamics are not significantly different. Several factors could explain this discrepancy. Timing may be specific to particular off-target loci, concentrations of Cas9 used in different experiments may influence enzyme availability for rarer off-targets, or the discrepancy in timescales between *in vitro* experiments (minutes) and *in vivo* experiments (hours to days) may push reactions toward equilibrium. So far, our data suggest that Cas9 does not temporally distinguish between on and off-targets in cells.

*In vitro* assays such as CIRCLE-Seq (*6*) and SITE-Seq (*7*) strip the DNA of all bound proteins before nuclease digestion, resulting in efficient cleavage of all potential sites by the Cas9 enzyme. These assays can detect potential off-targets with incredible sensitivity, but cannot provide information about how *in vitro* cleaved sites translate into *bona-fide* off-targets in cellular models. Chromatin state, modifications, other DNA-binding proteins, and collision with transcriptional complexes all influence Cas9 cutting efficiency and thus off-target mutagenesis *in vivo* (*14*, *30*, *46*). It is currently difficult to predict repair outcomes from *in vitro* data. Performing *in vitro* assays and then validating all potential cleavage sites by amplicon-NGS *in vivo* is subject to high false positive rates and can be extremely laborious depending on the number of potential sites, particularly when a single CIRCLE-Seq experiment can return as many as 3000 putative off-targets (*11*). DISCOVER-Seq instead directly provides data on true Cas9 cleavage events, as it measures off-targets within the edited cell in its original chromatin state. We have also found that DISCOVER-Seq robustly identifies true *in vivo* off-targets that are missed by CIRCLE-Seq or GUIDE-Seq due to poor ranking among hundreds to thousands of false positives. DISCOVER-Seq enables one to determine *bona-fide* off-targets in a one-step procedure, which reduces the number of target sites to be validated by amplicon-NGS by orders of magnitude. The *in vivo* sensitivity of DISCOVER-Seq is not as high as *in vitro* CIRCLE-Seq, as off-targets below 0.8% indels were missed. However, this could be due to the sensitivity of ChIP-Seq in tissues and might be potentially improved by using more sensitive ChIP methods such as CUT&RUN (*47*) or by increasing the overall read depth of the Illumina Sequencing.

MRE11 ChIP-Seq yields provocative data regarding the error-prone nature of Cas9 break repair. While indels formed by Cas9 involve error-prone repair, it is currently unclear to what extent these outcomes are the end-product of multiple rounds of error-free repair after genomic cleavage (*16*, *45*, *48*). Some gRNAs that exhibit low indel efficiency may actually cut efficiently but breaks could be repaired perfectly. This would be a complicating factor given that such sites would be overlooked when considering possible genomic translocations (*49*). DISCOVER-Seq detects off-target events prior to repair and is thus independent of the repair outcome. In the future, DISCOVER-Seq could also be used to find sites of Cas9 action that could be hot spots for translocations, rearrangements and larger deletions that do not result in small indels at the cut site. While DISCOVER-Seq signal is generally correlated with indel formation (**Fig. S4A**), there are notable exceptions. We found several target sites that showed disproportionately high MRE11-bound DNA fragments with low indels. For example the *D6Ertd527e* off-target in gP-edited B16-F10 cells exhibited a larger MRE11 peak than the on-target *Pcsk9* site, the off-target had only ∼25% indels compared to ∼66% indels at the on-target. More work remains to be done to determine if MRE11 binding can truly find sites that are perfectly repaired at high frequency, or if there are unknown limits to the predictive power between MRE11 binding and indel formation.

In summary, DISCOVER-Seq is broadly applicable to detect off-targets from all tested Cas variants in multiple species. It should also be applicable to other classes of nucleases such as Transcription activator-like effector nucleases (TALENs) and Zinc finger nucleases (ZFNs), though we have not tested these explicitly. The biological function of the MRN complex is highly conserved among species and even across domains and one could imagine applying a similar off-target detection strategy in plants or other organisms. Moreover, DISCOVER-Seq’s ability to visualize Cas enzyme activity in a cellular context has numerous applications beyond off-target discovery, including understanding CRISPR-Cas cutting patterns and dynamics *in vivo* and shedding light onto DSBs and repair outcomes in different cell types and tissues. DISCOVER-Seq can even be used to characterize clinical editing reagents and, in particular, to identify off-targets from personal genotypes uncovering how normal genetic variation affects editing accuracy. We also anticipate that it could potentially be used for real-time off-target discovery in patient biopsy samples after *in vivo* editing in the clinic.

## Acknowledgements

J.E.C., B.W., C.D.R., S.K.W., C.D.Y. and K.R.K. are supported by the Li Ka Shing Foundation and Heritage Medical Research Institute. J.E.C. is supported by the National Heart, Lung, and Blood Institute of the NIH under DP2-HL-141006. J.T.V. is supported by CIRM TRAN1-09292. B.W. is supported by a Sir Keith Murdoch Fellowship from the American Australian Association. This work used the Vincent J. Coates Genomics Sequencing Laboratory at UC Berkeley, supported by NIH S10 OD018174 Instrumentation Grant. We would like to thank Marie Johansson for technical support on intravenous tail vein injections and the Gladstone Stem Cell Core for their services.

## Contributions

B.W., S.K.W., C.D.R. and J.E.C. conceived and designed the study and experiments. B.W. performed ChIP-Seq, PCRs for amplicon-NGS and Western Blots in cell lines. S.K.W. analyzed ChIP-Seq and amplicon-NGS data and developed BLENDER software. C.D.Y. performed ChIP-qPCR time course experiments. K.R.K. prepared ChIP-Seq libraries. J.V.T. prepared amplicon-NGS libraries. P.A., M.J.P. and M. Morlock executed intravenous tail vein injections, animal terminations and organ collection and performed Western Blots in mouse tissues. L.M.J. and H.L.W. planned and performed iPSC editing experiments. M. Maresca and B.R.C. supervised experiments and provided expertise. J.E.C. supervised the study. B.W., S.K.W. and J.E.C wrote the manuscript with input from all authors.

## References

1. W.-J. Dai et al., CRISPR-Cas9 for in vivo Gene Therapy: Promise and Hurdles. Mol. Ther. Nucleic Acids. 5, e349 (2016).

2. S. Bae, J. Park, J.-S. Kim, Cas-OFFinder: a fast and versatile algorithm that searches for potential off-target sites of Cas9 RNA-guided endonucleases. Bioinformatics. 30, 1473–1475 (2014).

3. S. Abadi, W. X. Yan, D. Amar, I. Mayrose, A machine learning approach for predicting CRISPR-Cas9 cleavage efficiencies and patterns underlying its mechanism of action. PLoS Comput. Biol. 13, e1005807 (2017).

4. J.-P. Concordet, M. Haeussler, CRISPOR: intuitive guide selection for CRISPR/Cas9 genome editing experiments and screens. Nucleic Acids Res. 46, W242–W245 (2018).

5. D. Kim et al., Digenome-seq: genome-wide profiling of CRISPR-Cas9 off-target effects in human cells. Nat. Methods. 12, 237–43, 1 p following 243 (2015).

6. S. Q. Tsai et al., CIRCLE-seq: a highly sensitive in vitro screen for genome-wide CRISPRCas9 nuclease off-targets. Nat. Methods. 14, 607–614 (2017).

7. P. Cameron et al., Mapping the genomic landscape of CRISPR-Cas9 cleavage. Nat. Methods. 14, 600–606 (2017).

8. S. Q. Tsai et al., GUIDE-seq enables genome-wide profiling of off-target cleavage by CRISPR-Cas nucleases. Nat. Biotechnol. 33, 187–197 (2015).

9. W. X. Yan et al., BLISS is a versatile and quantitative method for genome-wide profiling of DNA double-strand breaks. Nat Commun. 8, 15058 (2017).

10. J. Lin, K.-C. Wong, Off-target predictions in CRISPR-Cas9 gene editing using deep learning. Bioinformatics. 34, i656–i663 (2018).

11. P. Akcakaya et al., In vivo CRISPR editing with no detectable genome-wide off-target mutations. Nature. 561, 416–419 (2018).

12. S. Kim, D. Kim, S. W. Cho, J. Kim, J.-S. Kim, Highly efficient RNA-guided genome editing in human cells via delivery of purified Cas9 ribonucleoproteins. Genome Res. 24, 1012–1019 (2014).

13. D. Kim et al., Genome-wide analysis reveals specificities of Cpf1 endonucleases in human cells. Nat. Biotechnol. 34, 863–868 (2016).

14. E. M. Kallimasioti-Pazi et al., Heterochromatin delays CRISPR-Cas9 mutagenesis but does not influence repair outcome. BioRxiv (2018), doi:10.1101/267690.

15. R. M. Yarrington, S. Verma, S. Schwartz, J. K. Trautman, D. Carroll, Nucleosomes inhibit target cleavage by CRISPR-Cas9 in vivo. Proc. Natl. Acad. Sci. USA. 115, 9351–9358 (2018).

16. C. D. Richardson, G. J. Ray, N. L. Bray, J. E. Corn, Non-homologous DNA increases gene disruption efficiency by altering DNA repair outcomes. Nat Commun. 7, 12463 (2016).

17. H. O’Geen, I. M. Henry, M. S. Bhakta, J. F. Meckler, D. J. Segal, A genome-wide analysis of Cas9 binding specificity using ChIP-seq and targeted sequence capture. Nucleic Acids Res. 43, 3389–3404 (2015).

18. X. Wu et al., Genome-wide binding of the CRISPR endonuclease Cas9 in mammalian cells. Nat. Biotechnol. 32, 670–676 (2014).

19. S. C. Knight et al., Dynamics of CRISPR-Cas9 genome interrogation in living cells. Science (80-.). 350, 823–826 (2015).

20. H. Ma et al., CRISPR-Cas9 nuclear dynamics and target recognition in living cells. J. Cell Biol. 214, 529–537 (2016).

21. R. Aleksandrov et al., Protein dynamics in complex DNA lesions. Mol. Cell. 69, 1046–1061.e5 (2018).

22. R. Shroff et al., Distribution and dynamics of chromatin modification induced by a defined DNA double-strand break. Curr. Biol. 14, 1703–1711 (2004).

23. J. S. Iacovoni et al., High-resolution profiling of gammaH2AX around DNA double strand breaks in the mammalian genome. EMBO J. 29, 1446–1457 (2010).

24. R. S. Williams, J. S. Williams, J. A. Tainer, Mre11-Rad50-Nbs1 is a keystone complex connecting DNA repair machinery, double-strand break signaling, and the chromatin template. Biochem Cell Biol. 85, 509–520 (2007).

25. J. Kobayashi, Molecular mechanism of the recruitment of NBS1/hMRE11/hRAD50 complex to DNA double-strand breaks: NBS1 binds to gamma-H2AX through FHA/BRCT domain. J Radiat Res. 45, 473–478 (2004).

26. C. D. Richardson et al., CRISPR-Cas9 genome editing in human cells occurs via the Fanconi anemia pathway. Nat. Genet. 50, 1132–1139 (2018).

27. T. J. Cradick, E. J. Fine, C. J. Antico, G. Bao, CRISPR/Cas9 systems targeting β-globin and CCR5 genes have substantial off-target activity. Nucleic Acids Res. 41, 9584–9592 (2013).

28. M. A. DeWitt et al., Sci Transl Med, in press, doi:10.1126/scitranslmed.aaf9336.

29. C. D. Richardson, G. J. Ray, M. A. DeWitt, G. L. Curie, J. E. Corn, Enhancing homology-directed genome editing by catalytically active and inactive CRISPR-Cas9 using asymmetric donor DNA. Nat. Biotechnol. 34, 339–344 (2016).

30. R. Clarke et al., Enhanced Bacterial Immunity and Mammalian Genome Editing via RNA-Polymerase-Mediated Dislodging of Cas9 from Double-Strand DNA Breaks. Mol. Cell. 71, 42–55.e8 (2018).

31. M. Jinek et al., A programmable dual-RNA-guided DNA endonuclease in adaptive bacterial immunity. Science (80-.). 337, 816–821 (2012).

32. M. W. Shen et al., Predictable and precise template-free CRISPR editing of pathogenic variants. Nature (2018), doi:10.1038/s41586-018-0686-x.

33. M. van Overbeek et al., DNA Repair Profiling Reveals Nonrandom Outcomes at Cas9-Mediated Breaks. Mol. Cell. 63, 633–646 (2016).

34. A. Taheri-Ghahfarokhi et al., Decoding non-random mutational signatures at Cas9 targeted sites. Nucleic Acids Res. 46, 8417–8434 (2018).

35. B. Zetsche et al., Cpf1 is a single RNA-guided endonuclease of a class 2 CRISPR-Cas system. Cell. 163, 759–771 (2015).

36. E. A. Boyle et al., High-throughput biochemical profiling reveals sequence determinants of dCas9 off-target binding and unbinding. Proc. Natl. Acad. Sci. USA. 114, 5461–5466 (2017).

37. B. P. Kleinstiver et al., High-fidelity CRISPR-Cas9 nucleases with no detectable genome-wide off-target effects. Nature. 529, 490–495 (2016).

38. C. A. Vakulskas et al., A high-fidelity Cas9 mutant delivered as a ribonucleoprotein complex enables efficient gene editing in human hematopoietic stem and progenitor cells. Nat. Med. 24, 1216–1224 (2018).

39. I. M. Slaymaker et al., Rationally engineered Cas9 nucleases with improved specificity. Science (80-.). 351, 84–88 (2016).

40. O. V. Evgrafov et al., Mutant small heat-shock protein 27 causes axonal Charcot-Marie-Tooth disease and distal hereditary motor neuropathy. Nat. Genet. 36, 602–606 (2004).

41. T. Geuens et al., Mutant HSPB1 causes loss of translational repression by binding to PCBP1, an RNA binding protein with a possible role in neurodegenerative disease. Acta Neuropathol. Commun. 5, 5 (2017).

42. L. Huang, J.-N. Min, S. Masters, N. F. Mivechi, D. Moskophidis, Insights into function and regulation of small heat shock protein 25 (HSPB1) in a mouse model with targeted gene disruption. Genesis. 45, 487–501 (2007).

43. S. H. Sternberg, S. Redding, M. Jinek, E. C. Greene, J. A. Doudna, DNA interrogation by the CRISPR RNA-guided endonuclease Cas9. Nature. 507, 62–67 (2014).

44. M. Shibata et al., Real-space and real-time dynamics of CRISPR-Cas9 visualized by high-speed atomic force microscopy. Nat Commun. 8, 1430 (2017).

45. E. K. Brinkman et al., Kinetics and Fidelity of the Repair of Cas9-Induced Double-Strand DNA Breaks. Mol. Cell. 70, 801–813.e6 (2018).

46. S. A. Verkuijl, M. G. Rots, The influence of eukaryotic chromatin state on CRISPR-Cas9 editing efficiencies. Curr. Opin. Biotechnol. 55, 68–73 (2018).

47. P. J. Skene, S. Henikoff, An efficient targeted nuclease strategy for high-resolution mapping of DNA binding sites. Elife. 6 (2017), doi:10.7554/eLife.21856.

48. L. Deriano, D. B. Roth, Modernizing the nonhomologous end-joining repertoire: alternative and classical NHEJ share the stage. Annu. Rev. Genet. 47, 433–455 (2013).

49. M. Kosicki, K. Tomberg, A. Bradley, Repair of double-strand breaks induced by CRISPRCas9 leads to large deletions and complex rearrangements. Nat. Biotechnol. 36, 765–771 (2018).

